# Neurons learn by predicting future activity

**DOI:** 10.1101/2020.09.25.314211

**Authors:** Artur Luczak, Bruce L. McNaughton, Yoshimasa Kubo

**Affiliations:** Canadian Center for Behavioural Neuroscience; University of Lethbridge, AB, Canada; Center for Neurobiology of Learning and Memory, University of California Irvine, USA

**Keywords:** Synaptic plasticity model, brain-inspired learning algorithm, neuronal population recordings

## Abstract

Understanding how the brain learns may lead to machines with human-like intellectual capacities. However, learning mechanisms in the brain are still not well understood. Here we demonstrate that the ability of a neuron to predict its future activity may provide an effective mechanism for learning in the brain. We show that comparing a neuron’s predicted activity with the actual activity provides a useful learning signal for modifying synaptic weights. Interestingly, this predictive learning rule can be derived from a metabolic principle, where neurons need to minimize their own synaptic activity (cost), while maximizing their impact on local blood supply by recruiting other neurons. This reveals an unexpected connection that learning in neural networks could result from simply maximizing the energy balance by each neuron. We show how this mathematically derived learning rule can provide a theoretical connection between diverse types of brain-inspired algorithms, such as: Hebb’s rule, BCM theory, temporal difference learning and predictive coding. Thus, this may offer a step toward development of a general theory of neuronal learning. We validated this predictive learning rule in neural network simulations and in data recorded from awake animals. We found that in the sensory cortex it is indeed possible to predict a neuron’s activity ∼10-20ms into the future. Moreover, in response to stimuli, cortical neurons changed their firing rate to minimize surprise: i.e. the difference between actual and expected activity, as predicted by our model. Our results also suggest that spontaneous brain activity provides “training data” for neurons to learn to predict cortical dynamics. Thus, this work demonstrates that the ability of a neuron to predict its future inputs could be an important missing element to understand computation in the brain.

## Introduction

Neuroscience is at the stage biology was at before Darwin. It has a myriad of detailed observations but no single theory explaining the connections between all of those observations. We do not even know if such a brain theory should be at the molecular level, or at the level of brain regions, or at any scale between. However, looking at Deep Neural Networks, which have achieved remarkable results in tasks ranging from cancer detection to self-driving cars, may provide useful insights. Although such networks may have different inputs and architectures, most of their impressive behavior can be understood in terms of the underlying common learning algorithm, called backpropagation (*1*).

Therefore, a better understanding of the learning algorithm(s) used by the brain could be central to develop a unifying theory of the brain function. There are two main approaches to investigate learning mechanisms in the brain: 1) experimentally, where persistent changes in neuronal activity are induced by a specific intervention (*2*); and 2) computationally, where algorithms are developed to achieve specific computational objectives while still satisfying selected biological constraints (*3, 4*). Here, we explore a different approach: 3) theoretical derivation, where a learning rule is derived from basic cellular principles, i.e. from maximizing metabolic energy of a cell. Using this approach, we found that maximizing energy balance by a neuron leads to a predictive learning rule, where a neuron adjusts its synaptic weights to minimize surprise: i.e. the difference between actual and predicted activity. Interestingly, this derived learning rule has a direct relation to some of the most promising biologically inspired learning algorithms, like predictive coding and temporal difference learning (see below), and Hebbian based rules can be seen as a special case of our predictive learning rule (see Discussion). Thus, our approach may provide a theoretical connection between multiple brain-inspired algorithms, and may offer a step toward development of a unified theory of neuronal learning.

There are multiple lines of evidence suggesting that the brain operates as a predictive system (*5-10*). However, it remains controversial as to how exactly predictive coding could be implemented in the brain (*4*). Most of proposed mechanisms involve specially designed neuronal circuits with “error units” to allow for comparing expected and actual activity (*11-14*). Although such models are motivated by cortical circuits, they require a precise network configuration, which could be difficult to achieve, considering number and variability of connections between each cortical neuron (*15, 16*). Moreover, many animal groups show an intelligent behavior even without cerebral cortex (e.g. octopus, crows). Thus, it is unlikely that such different brains share a common predictive circuit architecture. However, many basic properties of neurons are highly conserved throughout evolution (*17-19*). Therefore, we suggest that a single neuron using predictive learning rule could provide an elementary unit from which a variety of predictive brains may be built.

Interestingly, our predictive learning rule can also be obtained by modifying a temporal difference learning algorithm to be more biologically plausible. Temporal difference learning is one of the most promising ideas of how backpropagation-like algorithms could be implemented in the brain. It is based on using differences in neuronal activity to approximate top-down error signals (*4, 20-26*). A typical example of such algorithms is Contrastive Hebbian Learning (*27-29*), which was proven to be equivalent to backpropagation under certain assumptions (*30*). Contrastive Hebbian Learning requires networks to have reciprocal connections between hidden and output layers, which allows activity to propagate in both directions (Fig. 1A). The learning consists of two separate phases. First, in the ‘free phase’, a sample stimulus is continuously presented to the input layer and the activity propagates through the network until the dynamics converge to an equilibrium (activity of each neuron achieves steady-state level). In the second ‘clamped phase’, in addition to presenting stimulus to the input, the output neurons are also held clamped at values representing stimulus category (e.g.: 0 or 1), and the network is again allowed to converge to an equilibrium. For each neuron, the difference between activity in the clamped 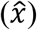 and free 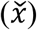 phases is used to modify synaptic weights (*w*) according to the equation (Eq. 1): 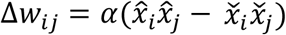, where *i* and *j* are indices of pre- and post-synaptic neurons respectively, and α is a small number representing learning rate. Intuitively, this can be seen as adjusting weights to push each neuron’s activity in the free phase, closer to the desired activity represented by the clamped phase. The obvious biological plausibility issue with this algorithm is that it requires the neuron to experience exactly the same stimulus twice in two separate phases, and that the neuron needs to ‘remember’ its activity from the previous phase. Our predictive learning rule provides a solution to this problem by predicting free phase steady-state activity, thus eliminating requirement for two separate stimulus presentations.

**Fig. 1.**
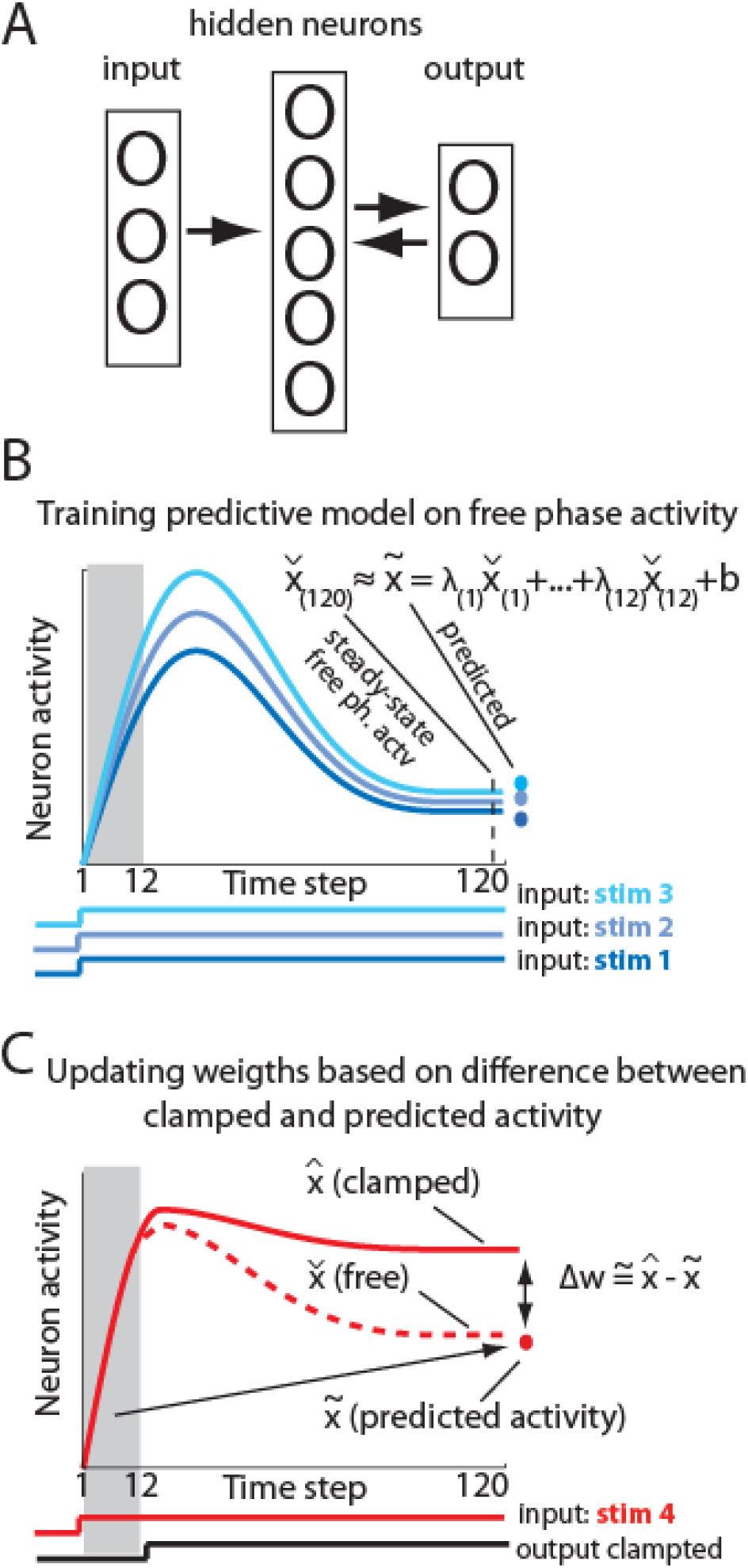
Basics of the algorithm. (A) Schematic of the network. Note that activity propagates back-and-forth between hidden and output layers. (B) Sample neuron activity in the free phase in response to different stimuli (marked with shades of blue). The free phase responses were used to train a linear model to predict a steady-state activity from the activity at earlier time steps (marked by shaded area; see main text for details). Bottom traces show duration of inputs, and dots represent predicted activity. (C) Activity of a neuron in response to a new stimulus with the network output clamped. Initially the network receives only input signal (free phase), but after a few steps the output signal is also presented (clamped phase; bottom trace). The red dot represents steady-state free phase activity predicted from initial activity (shaded region). For comparison, the dashed line shows a neuron’s activity in the free phase if the output is not clamped. Synaptic weights (*w*) are adjusted in proportion to the difference between steady-state activity in clamped phase 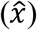 and predicted free phase activity 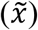.

For manuscript clarity, first we will describe how our predictive learning rule can be obtained by modifying Contrastive Hebbian Learning algorithm. Next, we will validate the predictive learning rule in simulation and in data recorded from awake animals, and we will show how our results shed new light on the function of spontaneous activity. The details of derivation of the learning rule by maximizing neuron energy balance will be presented at the end.

## Results

### Connection between predictive learning rule and Contrastive Hebbian Learning

As mentioned earlier, Contrastive Hebbian Learning algorithm requires network to converge to steady-state equilibrium in two separate learning phases, thus exactly the same stimulus has to be presented twice. However, this is unlikely to be the case in the actual brain. Here, we propose to solve this problem by combining both activity phases into one, which is inspired by sensory processing in the cortex. For example, in visual areas, when presented with a new picture, there is initially bottom-up driven activity containing mostly visual attributes of the stimulus (e.g. contours). This is then followed by top-down modulation containing more abstract information, e.g., “this object is a member of category x”, or “this object is novel” (Suppl. Fig. 1). Accordingly, our algorithm first runs only the initial part of the free phase, which represents bottom-up stimulus driven activity, and then, after a few steps the network output is clamped, corresponding to top-down modulation.

The novel insight here is that the initial bottom-up activity is enough to allow neurons to predict the steady-state part of the free phase activity, and the mismatch between predicted free phase and clamped phase can then be used as a teaching signal. To implement this idea in our model, for each neuron, activity during twelve initial time steps of the free phase 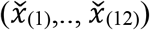 was used to predict its steady-state activity at time step 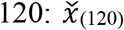 (Fig. 1B). Specifically, first, we presented sample stimuli in free phase to train linear model, such that: 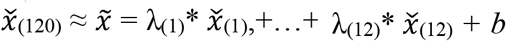, where 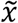 denote predicted activity, λ and *b* correspond to coefficients and offset terms of the least-squares model, and terms in brackets correspond to time steps. Next, a new set of stimuli was used for which free phase was ran only for the first 12 steps, and from step 13 the network output was clamped (Fig. 1C). The above least-squares model was then applied to predict free phase steady-state activity for each neuron, and the weights were updated based on the difference between predicted and clamped activity (Methods). Thus, to modify synaptic weights, in Eq. (1) we replaced activity in free phase with predicted activity 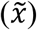: (Eq. 2) 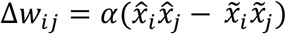). However, the problem is that this equation implies that a neuron needs also to know the predicted activity of all its presynaptic neurons 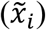, which could be not realistic. To solve this problem, we replaced 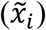 by the actual presynaptic activity in the clamped phase 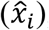, which we validated in network simulations (see the next section). This change leads to the following simplified synaptic plasticity rule (Eq. 3): 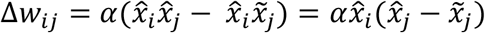. Thus, to modify synaptic weights, a neuron only compares its actual activity 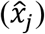 with its predicted activity 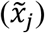, and applies this difference in proportion to each input contribution 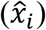.

### Predictive learning rule validation in neural network simulations

To test if predictive learning rule can be used to solve standard machine learning tasks, we created the following simulation. The neural network had 784 input units, 1000 hidden units, and 10 output units, and it was trained on a hand-written digit recognition task (MNIST (*31*); Suppl. Fig. 2; Methods). This network achieved 1.9% error rate, which is similar to neural networks with comparable architecture trained with the backpropagation algorithm (*31*). This demonstrates that the network with predictive learning rule can solve challenging non-linear classification tasks.

To verify that neurons could correctly predict future free phase activity, we took a closer look at sample neurons. Figure 2A illustrates the activity of all 10 output neurons in response to an image of a sample digit after the first epoch of training. During time steps 1-12, only the input signal was presented and the network was running in free phase. At time step 13, the output neurons were clamped, with activity of 9 neurons set to 0 and the activity of one neuron representing correct image class set to 1. For comparison, this figure also shows the activity of the same neurons without clamped outputs (free phase). It illustrates that, after about 50 steps in free phase, the network achieves steady-state, with predicted activity closely matching. When the network is fully trained, it still takes about 50 steps for the network dynamics in free phase to converge to steady-state (Fig. 2B). Note that, although all units initially increase activity at the beginning of the free phase, they later converge close to 0, except the one unit representing the correct category. Again, predictions made from the first 12 steps during free phase closely matched the actual steady-state activity. The hidden units also converged to steady-state after about 50 steps. Figure 2C illustrates the response of one representative hidden neuron to 5 sample stimuli. Because hidden units experience the clamped signal only indirectly, through synapses from output neurons, their steady-state activity is not bound to converge only to 0 or 1, as in the case of output neurons. Actual and predicted steady-state activity for hidden neurons is presented in Figure 2D. The average correlation coefficient between predicted and actual free phase activity was R=1±0.0001SD (averaged across 1000 hidden neurons in response to 200 randomly selected test images). Note that for all predictions we used a cross-validation approach, where we trained a predictive model for each neuron on a subset of the data and applied it to new examples, which were then used for updating weights (Methods). Thus, neurons were able successfully to generalize their predictions to new unseen stimuli. The network error rate for the training and test dataset is shown in Fig. 2E. This demonstrates that the predictive learning rule worked well, and each neuron accurately predicted its future activity.

**Fig. 2.**
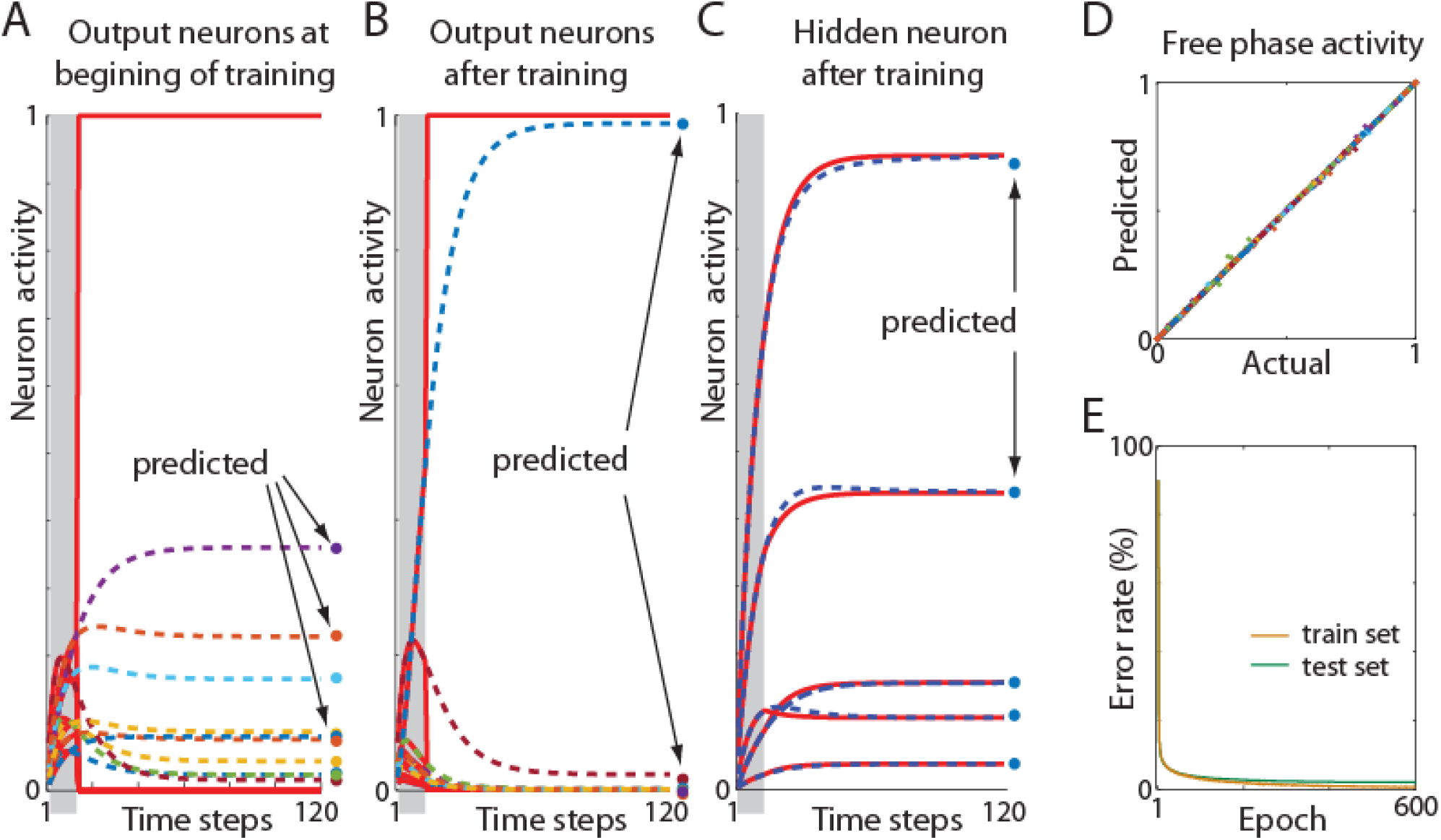
Neuron prediction of expected activity. (A) Activity of 10 output neurons in response to a sample stimulus at the beginning of the network training. Gray area indicates the extent of the free phase (time steps 1-12). Solid red lines show activity of the neurons clamped at step 13. For comparison, dashed lines represent free phase activity if output neurons had not been clamped. Dots show predicted steady-state activity in free phase based on initial activity (from steps 1-12). (B) Activity of the same neurons after network training. Note that free phase and predicted activity converged to desired clamped activity. (C) Activity of a representative neuron in the hidden layer in response to 5 different stimuli after network training. Solid and dashed lines represent clamped and free phase respectively, and dots show predicted activity. (D) Predicted vs. actual free phase activity. For visualization clarity only every tenth hidden neuron out of 1000 is shown, in response to 20 sample images. Different colors represent different neurons, but some neurons may share the same color due to the limited number of colors. Distribution of points along the diagonal shows that predictions are accurate. (E) Decrease in error rate across training epochs. Yellow and green lines denote learning curves for training and test data set respectively. Note that in each epoch we only used 2% out of 60,000 training examples.

### Biologically motivated network architectures

We also tested the predictive learning rule in multiple other network architectures, which were designed to reflect additional aspects of biological neuronal networks. First, we introduced a constraint that 80% of the hidden neurons were excitatory, and the remaining 20% had only inhibitory outputs. This follows observations that biological neurons release either excitatory or inhibitory neurotransmitters, not both (Dale’s law (*32*)), and that about 80% of cortical neurons are excitatory. The network with this architecture achieved an error rate of 2.66% (Suppl. Fig. 3A). We also tested our algorithm in a network without symmetric weights, which resulted in similar performance as the original network (1.96%, Suppl. Fig. 3B). Moreover, we implemented the predictive learning rule in a network with spiking neurons, which again achieved a similar error rate of 2.46% (Suppl. Fig. 4). Our predictive learning rule was also tested in a deep convolutional network (Fig. 3A), which architecture was shown to resemble neuronal processing in the visual system (*33, 34*). Using this convolutional network, we tested our algorithm on a more challenging dataset for biologically inspired algorithms: CIFAR-10 (*35*). This dataset consists of color images representing 10 different classes like: airplanes, cars, birds, cats, etc. We achieved 20.03% error rate, which was comparable with training the same network using backpropagation through time algorithm (Fig. 3B; see Methods for details; code to reproduce those results is available at: https://github.com/ykubo82/bioCHL/tree/master/conv). Altogether, this shows that our predictive learning rule performs well in a variety of biologically motivated network architectures.

**Fig. 3.**
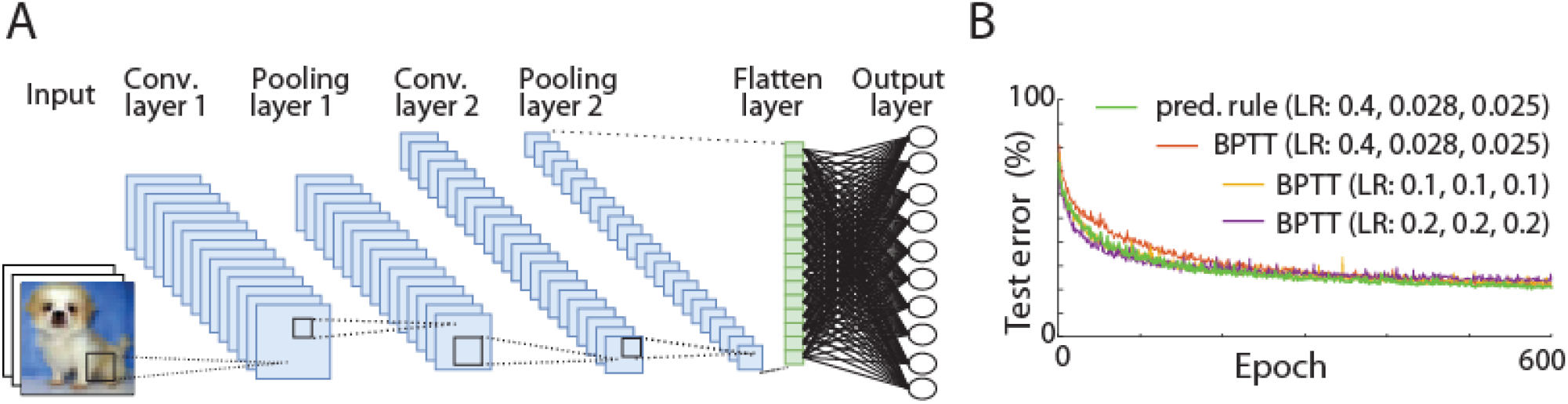
Implementation of predictive learning rule in a multilayer convolutional neuronal network. (A) Depiction of our convolutional network architecture (see Methods). (B) Learning curves for convolutional network trained using a predictive learning rule (green) and for comparison the same network trained using backpropagation through time (BPTT). The red line shows a learning curve for BPTT using the same learning rates as in our predictive model (RL: 0.4, 0.028, 0.025); yellow line: BPTT with learning rate of 0.1 for all layers, and violet line: BPTT with learning rate of 0.2 for all layers. It illustrates that on CIFAR-10, performance of the deep network using our predictive learning rule was comparable with BPTT.

### Predictive learning rule validation in awake animals

To test whether real neurons could also predict their future activity, we analyzed neuronal recordings from the auditory cortex in awake rats (Methods). As stimuli we presented 6 tones, each 1s long and interspersed by 1s of silence, repeated continuously for over 20 minutes (Suppl. Materials). For each of the 6 tones we calculated separately average onset and offset response, giving us 12 different activity profiles for each neuron (Fig. 4A). For each stimulus, the activity in the time window 15-25ms was used to predict average future activity within the 30-40ms window. We used 12-fold cross-validation, where responses from 11 stimuli were used to train the least-square model, which was then applied to predict neuron activity for the 1 remaining stimulus. This procedure was repeated 12 times for each neuron. The average correlation coefficient between actual and predicted activity was R = 0.36±0.05 SEM (averaged across 55 cells from 4 animals, Fig. 4B). Distribution of correlations coefficients for individual neurons were significantly different from 0 (t-test p<0.0001; insert in Fig. 4B). This shows that neurons have predictable dynamics, and from an initial neuronal response its future activity could be estimated.

**Fig. 4.**
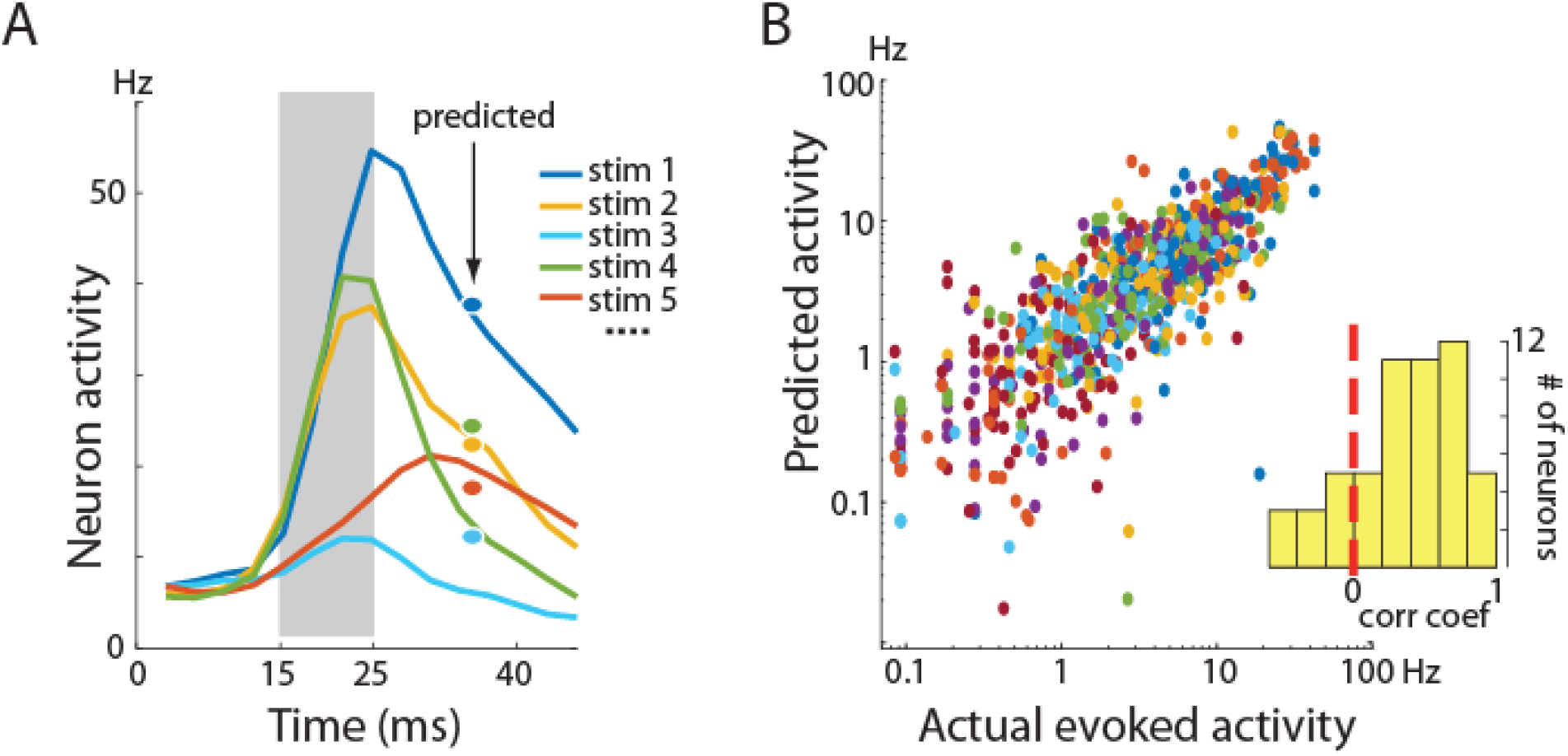
Predicting future activity of cortical neurons. (A) Response of a representative neuron to different stimuli. For visualization only 5 out of 12 responses are shown. Gray area indicates the time window which was used to predict future activity. Dots show predicted average activity in the 30-40ms time window. Colors correspond to different stimuli. (B) Actual vs. predicted activity for 55 cells from 4 animals in response to 12 stimuli. Different colors represent different neurons, but some neurons may share the same color due to the limited number of colors. Insert: histogram of correlation coefficients for individual neurons. Skewness of the distribution to the right shows that for most neurons the correlation between actual and predicted response was positive.

However, much stronger evidence supporting our learning rule is provided by predicting long-term changes in cortical activity. Specifically, repeated presentation of stimuli over tens of minutes induces long-term changes in neuronal firing rates (*36*), similar as in perceptual learning. Importantly, based on our model, it was possible to infer which individual neurons will increase or decrease their firing rate. To explain it, first let’s look at neural network simulations in Fig. 5A. It shows that, for a neuron, the average change in activity from one learning epoch to the next, depends on the difference between clamped (actual) activity and predicted (expected) activity, in the previous learning epoch (Fig. 5A; correlation coefficient R = 0.35, p<0.0001; Suppl. Materials). Similarly, for cortical neurons, we found that the change in firing rate from the 1st to the 2nd half of the experiment was positively correlated with differences between evoked and predicted activity during the 1st half of experiment (R = 0.58, p<0.0001; Fig. 5B, Suppl. Materials). Those changes in activity patterns were blocked by an NMDA receptor antagonist, as we showed using this data in (*36*), which provides strong support that this phenomenon depends on synaptic plasticity. Results presented in Fig. 5 could be understood in terms of Eq. 3, where if actual activity is higher than predicted, then synaptic weights are increased, thus leading to higher activity of that neuron in the next epoch. Therefore, similar behavior of artificial and cortical neurons, where firing rate changes to minimize ‘surprise’: difference between actual and predicted activity, provides a strong evidence in support of the predictive learning rule presented here.

**Fig. 5.**
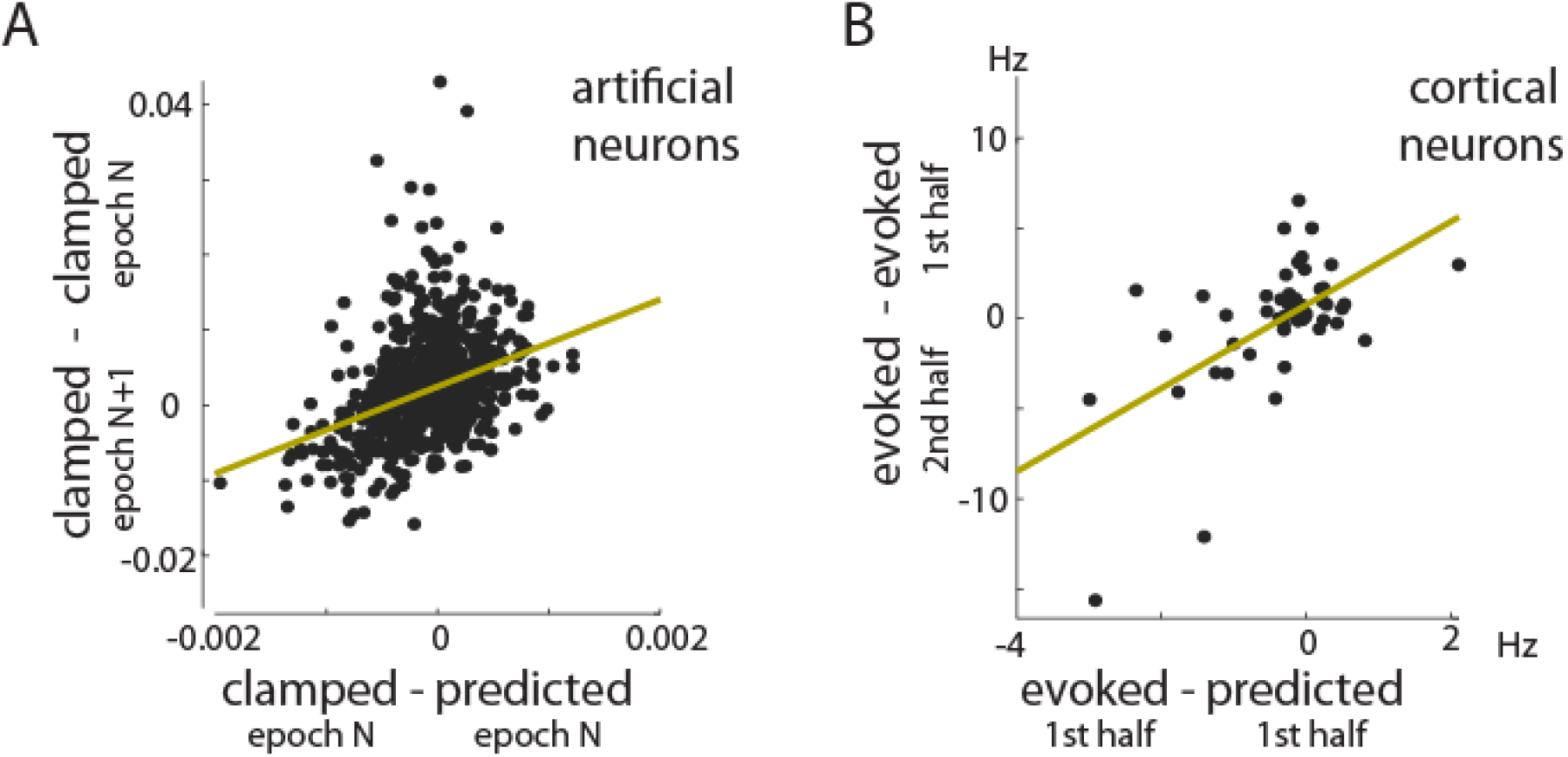
Long-term changes in neuronal activity in our model and in cortical neurons. (A) Average change in clamped steady-state activity between 2 consecutive learning epochs in our network model. This change relates to the difference between clamped and predicted activity in the earlier epoch (*N*=7; Suppl. Materials). Each dot represents one neuron. Regression line is shown in yellow. (B) Average change in firing rate between 1st and 2nd half of our experiment with repetitive auditory stimulation. This firing rate change correlates to the difference between stimulus evoked and predicted activity during the 1st half of the experiment (Suppl. Materials). Each dot represents the activity of one neuron averaged across stimuli. The similar behavior of cortical and artificial neurons suggests that both may be using essentially the same learning rule. Thus, this evidence that a neuronal change in firing rate relates to value of predicted activity, provides a novel insight about neuronal plasticity.

### Deriving predictive model parameters from spontaneous activity

Next, we tested whether spontaneous brain activity could also be used to predict neuronal dynamics during stimulus presentation. Spontaneous activity, such as during sleep, is defined as an activity not directly caused by any external stimuli. However, there are many similarities between spontaneous and stimulus evoked activity (*37-40*). For example, spontaneous activity is composed of ∼50-300 ms long population bursts called packets, which resemble stimulus evoked patterns (*41*). This is illustrated in Figure 6A, where spontaneous activity packets in the auditory cortex are visible before sound presentation (*42, 43*). In our experiments, each 1s long tone presentation was interspersed with 1s of silence, and the activity during 200-1000 ms after each tone was considered as spontaneous (animals were in soundproof chamber; Suppl. Materials). The individual spontaneous packets were extracted to estimate neuronal dynamics (Methods). Then the spontaneous packets were divided into 10 groups based on similarity in PCA space (Suppl. Materials), and for each neuron we calculated its average activity in each group (Fig. 6B). As in previous analyses in Fig 4A, the initial activity in time window 5-25ms was used to derive the least-square model to predict future spontaneous activity in the 30-40ms time window (Suppl. Materials). This least-square model was then applied to predict future evoked responses from initial evoked activity for all 12 stimuli. Figure 6C shows actual vs predicted evoked activity for all neurons and stimuli (correlation coefficient R = 0.2±0.05 SEM, averaged over 40 cells from 4 animals; the insert shows the distribution of correlation coefficients of individual neurons; p=0.0008, t-test). Spontaneous brain activity is estimated to account for over 90% of brain energy consumption (*44*), however the function of this activity still is a mystery. The foregoing results offer a new insight: because neuronal dynamics during spontaneous activity is similar to evoked activity, thus spontaneous activity can provide “training data” for neurons to build a predictive model.

**Fig. 6.**
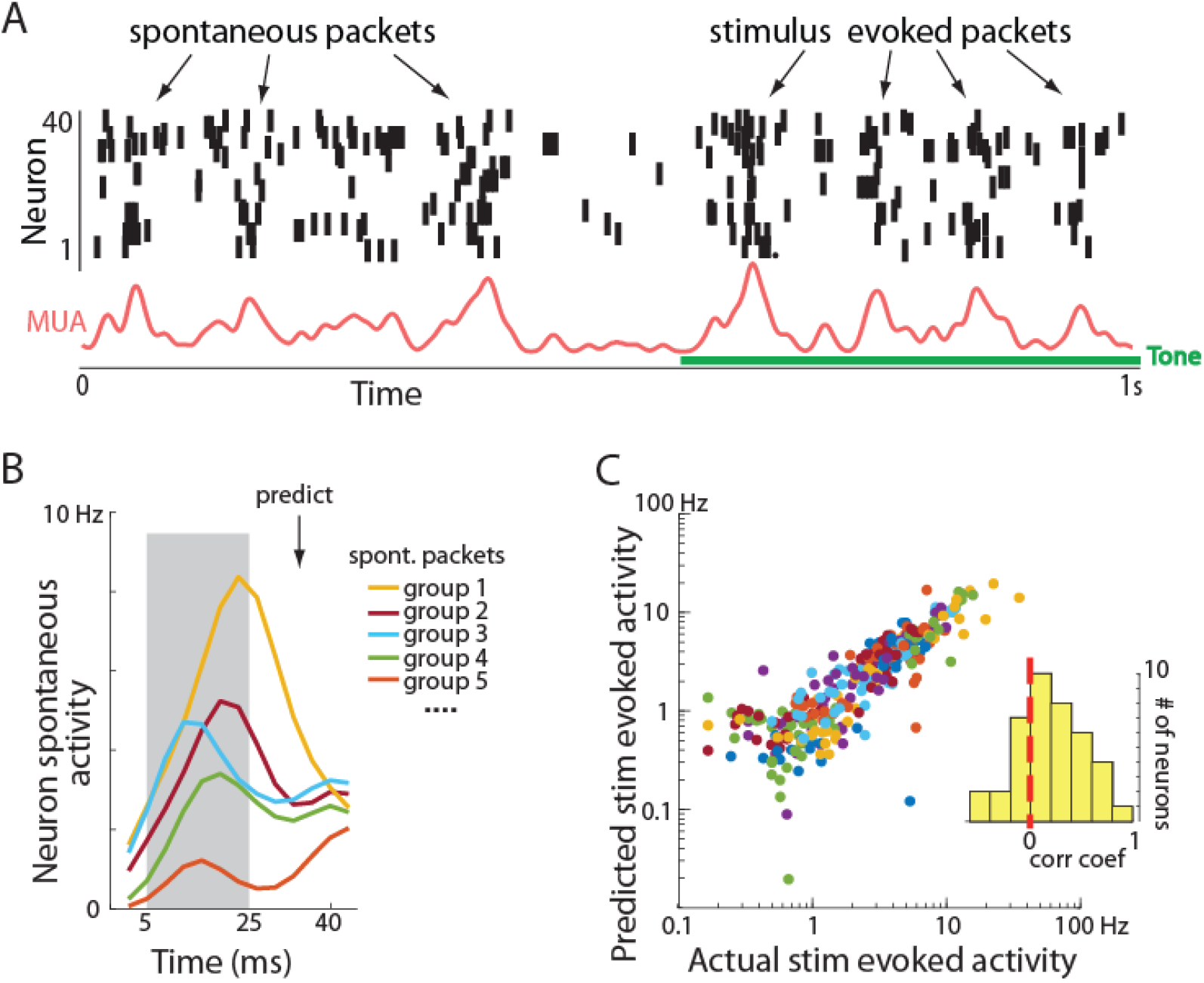
Predicting stimulus evoked responses from spontaneous activity dynamics. (A) Sample spiking activity in the auditory cortex before and during tone presentation. Note that spontaneous activity is not continuous, but rather composed of bursts called packets which are similar to tone evoked packets. The red trace shows smoothed multiunit activity: summed activity of all neurons (adopted with permission from (*43*)). (B) Spontaneous packets were divided into 10 groups based on population activity patterns. Activity of a single neuron in 5 different spontaneous packet groups is shown. Gray area indicates the time window used for predicting future average activity within the 30-40ms time window (marked by arrow). This predictive model derived from spontaneous activity was then applied to predict future evoked activity based on initial evoked response. (C) Actual vs. predicted tone evoked activity. Plot convention is the same as in Fig. 3B. Skewness of the histogram to the right shows that for most neurons the evoked dynamics can be estimated based on spontaneous neuron’s activity.

### Predictive learning rule derivation by maximizing neuron energy balance

Interestingly, predictive learning rule in Eq. 3: 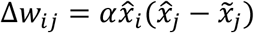 is not an *ad hoc* algorithm devised to solve a computational problem, but this form of learning rule arises naturally as a consequence of minimizing a metabolic cost by a neuron. Most of the energy consumed by a neuron is for electrical activity, with synaptic potentials accounting for ∼50% and action potential for ∼20% of ATP used (*45*). Using a simplified linear model of neuronal activity, this energy consumption for a neuron *j* can be expressed as 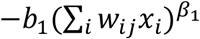, where *x*_*i*_ represents the activity of pre-synaptic neuron *i, w* represents synaptic weights, *b*_*1*_ is a constant to match energy units, and *β*_*1*_ describes a non-linear relation between neuron activity and energy usage, which is estimated to be between 1.7 - 4.8 (*46*). The remaining ∼30% of neuron energy is consumed on housekeeping functions, which could be represented by a constant −*ε*. On the other hand, the increase in neuronal population activity also increases local blood flow leading to more glucose and oxygen entering a neuron (see review on neurovascular coupling: (*47*)). This activity dependent energy supply can be expressed as: 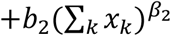, where *x*_*k*_ represents spiking activity of neuron *k* from a local population of *K* neurons (*k* ∈ {1, …, *j*, … *K*}); *b*_*2*_ is a constant and *β*_*2*_ reflects the exponential relation between activity and blood volume increase, which is estimated to be in range *β*_*2*_: 1.7-2.7 (*46*). Note that sum of local population activity ∑_*k*_ *x*_*k*_, also includes activity of neuron *j*: *x*_*j*_ = ∑_*i*_ *w*_*ij*_*x*_*i*_, as all local neurons contribute to local neurovascular coupling. Putting all the above terms together, the energy balance of a neuron *j* could be expressed as

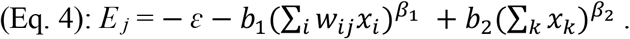

This formulation shows that to maximize energy balance, a neuron has to minimize its electrical activity (be active as little as possible), but at the same time, it should maximize its impact on other neurons’ activities to increase blood supply (be active as much as possible). Thus, weights have to be adjusted to strike a balance between two opposing demands: maximizing the neuron’s downstream impact and minimizing its own activity (cost). This energy objective of a cell could be paraphrased as *“lazy neuron principle”: maximum impact with minimum activity*.

We can calculate such required changes in synaptic weights Δ*w* that will maximize neuron’s energy *E* _*j*_ by using gradient ascent method. For that, we need to calculate derivative of *E* _*j*_ with respect to *w*_*ij*_

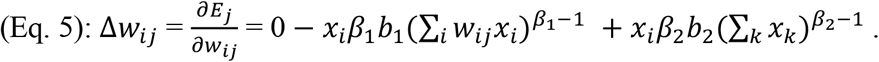

The appearance of *x*_*i*_ in the last term in (Eq. 5) comes from the fact that ∑_*k*_ *x*_*k*_, includes *x*_*j*_ which is function of *w*_*ij*_*x*_*i*_, as explained above. Thus, if we denote population activity as: 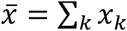, and considering that ∑_*i*_ *w*_*ij*_*x*_*i*_ = *x*_*j*_, then after moving *x*_*i*_ in front of brackets and after switching order of terms we obtain:

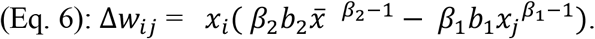

In case that *β*_*1*_ = 2 and *β*_*2*_ = 2, this formula simplifies from exponential to linear. However, even if *β*_*1*_ and *β*_*2*_ are anywhere in the range: 1.7 < *β*_*1*_ < 4.8 and 1.7 < *β*_*2*_ < 2.7 (*46*), the expression 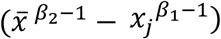 still is well approximated by its linearized version: 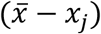 for typical values of *x* in range 0-1 (Suppl. Fig. 6). After also denoting that α_1_ = *β*_1_ *b*_1_ and 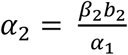 and after taking α_1_ in front of brackets, we obtain:

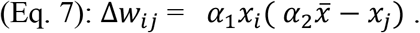

Although in this derivation we used linear model of a neuron, including a non-linear neural model like ReLU: *f*(*x*) = *x*^+^ = max(0, *x*) leads to similar expression (Suppl. Materials). Moreover, if we use the same derivation steps but to maximize neuron energy balance *in the future*, then Eq. 7, changes to

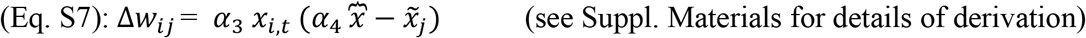

Note that Eq. S7 has the same form as the predictive learning rule in Eq. 3: 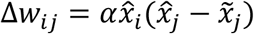. Here, 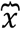 represents population recurrent activity, which can be thought of as top-down modulation, similar to 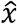. Also note that activity of neuron *j*: *x*_*j*_ from Eq.7, became here future predicted activity: 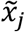. Thus, this derivation shows that the best strategy for a neuron to maximize future energy resources requires predicting its future activity. Altogether, this reveals an unexpected connection, that learning in neural networks could result from simply maximizing energy balance by each neuron.

## Discussion

Here we present theoretical, computational and biological evidence that the basic principle underlying single neuron learning may rely on minimizing future surprise: a difference between actual and predicted activity. Thus, a single neuron is not only performing summation of its inputs, but it also predicts the expected future, which we propose is a crucial component of the brain’s learning mechanism. Note that a single neuron has complexity similar to single cell organisms, which were shown to have ‘intelligent’ adaptive behaviors, including predicting consequences of its action in order to navigate toward food and away from danger (*48-50*). This suggests that typical neuronal models used in machine learning may be too simplistic to account for the essential computational properties of biological neurons. Our work suggests that a predictive mechanism may be an important computational element within neurons, which could be crucial to understand learning mechanisms in the brain.

This is supported by a theoretical derivation showing that the predictive learning rule provides an optimal strategy for maximizing metabolic energy of a neuron. To our knowledge, this is the first time where a synaptic learning rule has been derived from basic cellular principles, i.e. from maximizing energy of a cell. This provides a more solid theoretical basis over previous biologically-inspired algorithms, which were developed *ad hoc* to solve specific computational tasks while still satisfying selected biological constraints. However, it should be emphasized that many of those previous algorithms provided novel and insightful ideas which enabled the development of our model. Importantly, our derived learning rule provides a theoretical connection between those diverse types of brain-inspired algorithms, as discussed below.

One of the most influential ideas about brain’s learning algorithm was proposed by Donald Hebb, which is based on correlated firing: a.k.a. ‘cells that fire together wire together’ (*51*). This could be written as: Δ*w*_*ij*_ ∝ *x*_*i*_*x*_*j*_, where Δ*w*_*ij*_ is change in synaptic weight between neurons *i* and *j*, ∝ denotes proportionality, and *x*_*i*_ and *x*_*j*_ represents pre- and post-synaptic activity, respectively. Note that this is a special case of our predictive learning rule: 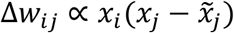 when 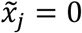, i.e. when a neuron does not make any prediction (note that here *x*_*i*_ and *x*_*j*_ represent actual activity as it is the case in clamped phase (i.e. 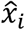 and 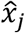 in Eq.3), thus for comparison clarity, hat symbol ^ can be omitted here). Despite its influential role, the original Hebb’s rule was shown to be unstable, as the synaptic weights will tend to increase or decrease exponentially. To solve this problem, a BMC theory was proposed (*52*), which can be expressed in a simplified form as: Δ*w*_*ij*_ ∝ *x*_*i*_(*x*_*j*_ − *θ*_*j*_)*x*_*j*_, where *θ*_*j*_ can be considered as the average activity of neuron *j* across all input patterns. Note that if in our equation: 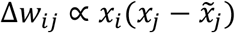, we would use the simplest predictive model: always predicting the average activity, then 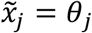 and our predictive rule becomes equivalent to the core part of BCM rule and could be seen as a linearized version of the full BCM rule. However, it was noted that networks trained using the BCM rule do not achieve the same level of accuracy as other learning rules (*53*). It is consistent with our experience that performance of our algorithm deteriorated when we used average activity of each neuron for predictions. We interpret from this, that dynamically adjusting predictions based on most recent activity allows for more precise weight adjustments.

Moreover, we described in the results section, how our predictive learning rule directly relates to Contrastive Hebbian Learning, which belongs to class of temporal difference learning algorithms. Our algorithm is also similar to other predictive algorithms (see Introduction). The main difference is that we propose that neurons can internally calculate their predictions, rather than relying on specialized neuronal circuits, as pointed out by the Reviewer. We already mentioned earlier that organisms with simpler neuronal systems may not have predictive circuits as proposed to exist in the cortex (*12, 14*). Thus, a predictive learning rule at the level of a single neuron may provide more basic description of the learning process across different brains. However, our model should not be taken as precluding possibility that in more complex brains, in addition to intracellular predictions, neurons may form predictive circuits to enhance predictive abilities of an organism. Our model is also closely related to the work of (*54-56*) where depolarization of basal dendrites serves as a prediction of top down signals from apical dendrites in pyramidal neurons. Again, our derived model could be seen as generalization of those ideas as it is not constrained to any specific cell type. The other interesting aspect of our model is that it belongs to the category of energy-based models, for which it was shown that synaptic update rules are consistent with spike-timing-dependent plasticity (*57*). Considering all the above, we suggest that our plasticity rule derived from basic metabolic principles could serve as a common denominator for diverse types of biologically inspired learning algorithms, and as such, it may offer a step toward development of a unified neuronal learning theory.

Biological neurons have a variety of cellular mechanisms which operate on time scales of 1∼100ms suitable for implementing predictions (*58-62*). The most likely mechanism appears to be calcium signaling. For example, when a neuron is activated, it leads to corresponding elevation of somatic calcium for tens of ms (*63*). This time period with elevated calcium could indicate that a certain level of new input is expected to arrive in that time window. For instance, if bottom-up visual stimulus triggers multiple spikes in a neuron, then the resulting proportional increase in calcium concentration may signal that a higher level of follow up activity is expected, which could correspond to predicting a higher level of e.g. top-down modulation. This would be consistent with our experimental data, where higher activity at stimulus onset is correlated with higher activity ∼20 ms later (Fig. 4). More direct evidence of calcium signaling involvement in predictive activity is provided by experiments where calcium plateau potentials are triggered by bursts of action potentials (complex spikes) in CA1 pyramidal cells. It was shown that such plateau potentials can result in a rapid synaptic plasticity to produce predictive place cell activity (*64*). There are also other possible cellular properties which could support predictive mechanisms. For instance, it was shown that neurons can preferentially respond to inputs arriving at specific resonance frequencies (range: ∼1-50 Hz) (*65, 66*). This provides another example that neurons do have cellular mechanisms to ‘remember’ and to ‘act’ accordingly based on their past activity tens of ms earlier (*60*). Interestingly, the core prediction of BCM and our model that synaptic weights should increase/decrease if neuron is stimulated above/below expected activity, is supported by experimental evidence where applying strong/weak electrical stimulation to cells in CAl induced LTP/LTD, respectively (*67*). All of this demonstrates compatibility of our model with neurophysiology. Unfortunately, due to limits in current technology, it could be challenging to experimentally probe subcellular mechanisms to directly test our predictive learning rule. However, considering the cellular mechanisms listed above and the consistency of our model with experimental data presented in Figs 4-6, altogether it shows that neurons are at least capable of implementing the predictive learning rule.

Our work also suggests that packets could be basic units of information processing in the brain. It is well established that sensory stimuli evoke coordinated bursts (packets) of neuronal activity lasting from tens to hundreds of ms. We call such population bursts packets, because they have stereotypical structure, with neurons active at the beginning conveying bottom-up sensory information (e.g. this is a face), and later in the packet neurons represent additional higher order information (e.g. this is a happy face of that particular friend)(*68*). Also the later part of the packet can encode if there is discrepancy with expectation (e.g. this is a novel stimulus (*69, 70*); Suppl. Fig. 1). This is likely because only the latter part of the packet can receive top-down modulation after information about that stimulus is exchanged between other brain areas, which is the case even during passive stimulus presentation (*71, 72*). Thus, our work suggests that the initial part of the packet can be used to infer what the rest of the brain may ‘think’ about this stimulus, and the difference from this expectation can be used as a learning mechanism to modify synaptic connections. This could be the reason why, for example, we cannot process visual information faster than ∼20 frames/s, as only after evaluating if a given image is consistent with expectation, can the next image be processed by the next packet, which takes ∼50ms. Our predictive learning rule thus implies, that sensory information is processed in discrete units and each packet may represent an elementary unit of perception.

When recording neuronal activity in the cortex, the slowest oscillations (<10Hz) are by far the most dominant (*43, 73*), and it is one of the biggest questions in neuroscience: what is the function of those oscillations (*74*). Therefore it is worth noticing how learning rule derived from the basic cellular principles may relate to packets which are the main part of slow oscillations (*41, 75, 76*). As described above, dividing information into discrete packets, could provide an effective mechanism to improve neuronal predictions. It could allow for easier differentiation of feed-forward signals arriving during the initial wave of a packet from predicted top-down information arriving later. Another big question in neuroscience is about the function of spontaneous brain activity (*44*). For example, why would the brain spend so much energy to generate packets even during e.g. sleep? Interestingly, similarly as in the brain where most energy is consumed by spontaneous activity (*44*), in our model most energy (i.e. computational time) is used for free phase network activity, which allows intracellular predictive model to learn network dynamics in an unsupervised way. Thus, free phase activity in our model suggests that the function of spontaneous packets could be to provide neurons with diverse training data to improve the robustness of the predictive model, as supported by results presented in Fig. 6. Moreover, note that free phase activity may also be used for unsupervised learning. For example, if new input is present in the free phase, neurons can still calculate if such evoked activity is consistent with internal model predictions. If not, then weights can be modified to get free phase activity evoked by new stimuli to be closer to the prediction (it is the same mechanism as we use in clamped phase during supervised learning). This is a similar idea to unsupervised pre-training (Hinton and Salakhutdinov, 2006), however more future work is needed to investigate it.

## Methods

### Neural Network (MNIST dataset)

The code for our network with predictive learning rule, which we used to produce results presented in Fig. 2 is available at https://github.com/ykubo82/bioCHL, which contains all implementation details. Briefly, the base network has the architecture: 784-1000-10 with sigmoidal units, and with symmetric connections (see Suppl. Fig. 3-4 for more biologically plausible network architectures which we also tested). Neuron activity dynamics in hidden layer is described as in standard network with Contrastive Hebbian Learning (*77*):

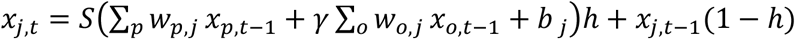

where *w*_*p,j*_ denotes weight from neuron *p* in input layer to neuron *j* in hidden layer, *w*_*o,j*_ denotes weight from output layer neuron to hidden layer neuron *j, b* is a bias, *t* is a time step and *S* is a sigmoid activation function. Parameter *h=0*.*1* is the Euler method’s time-step commonly used to improve computational stability. However, changing *h* to 0.2 or 1 resulted in similar network performance here. In the standard implementation of Contrastive Hebbian Learning, all top-down connections *w*_*o,j*_ are also multiplied by a small number *γ* (∼0.1) (*77*). This different treatment of feed-forward and feedback connections could be biologically questionable as many brain circuits are highly recurrent and e.g. granule cells do not seem to have specific dendrites for receiving feedback signals. Therefore, to make our network more biologically plausible we set this feedback gain factor *γ* to 1, thus allowing our network to learn by itself what should be the contribution of each input. For output layer, term ∑_*o*_ *w*_*o,j*_ *x*_*o,t*−1_ is set to 0 as there are no top-down connections to that layer. Neurons in the input layer do not have any dynamics as their activity is set to a value corresponding to pixel intensity in the presented image. To accelerate training, we used AdaGrad (*78*), and we applied a learning rate of 0.03 to the hidden layer and 0.02 for the output layer. Synaptic weights for neurons in hidden and output layers were modified as described in Eq. 3.

### Future activity prediction

For all the predictions we used a cross-validation approach. Specifically, in each training cycle we ran free phase on 490 examples, which were used to derive least-squares model for each neuron to predict its future activity at time step 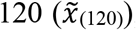, from its initial activity at steps 1-12 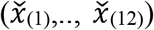. This can be expressed as: 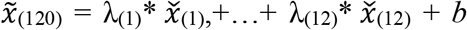, where terms in brackets correspond to time steps, and λ and *b* correspond to coefficients and offset terms found by least-squares method. Next, 10 new examples were taken, for which free phase was ran only for 12 steps, then the above derived least-squares model was applied to predict free phase steady-state activity for each of 10 examples. From step 13 the network output was clamped. The weights were updated based on the difference between predicted and clamped activity calculated only from those 10 new examples. This process was repeated 120 times in each training epoch. Moreover, the MNIST dataset has 60,000 examples which we used for the above described training, and 10,000 additional examples which were only used for testing. For all plots in Figure 2 we only used test examples which network never saw during training. This demonstrates that each neuron can accurately predict its future activity even for novel stimuli which were never presented before.

### Convolutional Neural Network (CIFAR-10 dataset)

The convolutional network has an input layer of size: 32×32×3, corresponding to size of a single image with 3 color channels in CIFAR-10 dataset (this dataset consists of 5000 training and 1000 test images for each of 10 classes (*35*)). The network has two convolutional and pooling layers followed by one fully connected output layer (Fig. 3A). The filter size for all the convolutional layers is 3×3 with stride 1, and the number of filters is 256 and 512 for first and second convolutional layers, respectively. We did not use zero-padding. For pooling, we used the max pooling with 2×2 filters and stride 2. The activation function for the convolutional and the fully connected layers was the hard-sigmoid activation function, *S(x) = (1+hardtanh(x-1))*0*.*5*, as implemented in (*26*). The learning rates were: 0.4, 0.028, and 0.025 for the first, second convolutional layer, and for the fully connected output layer, respectively. The Euler method’s time-step *h* was set to 1. Considering that clamping output neurons at only two extreme values: 0 or 1, may not be the most accurate model of top-down signals in the brain, thus here we implemented weak clamping as proposed in (*25*). Shortly, instead of setting the value of the output neuron to 0 or 1 during clamped phase, output neurons were only slightly nudged toward required values. For example, if an output neuron should have value of 1, then it was clamped at value: 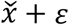, where 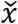 is free phase steady-state activity of that output neuron, and *ε* is a small nudging factor toward 1. To calculate nudging for each neuron we used a clamping factor of 0.01 as described in (*25*). This network with our predictive learning rule achieved 20.03% accuracy on CIFAR-10 dataset. Using the original ‘hard’ clamping, changing *h* to 0.1 or increasing number of neurons to 326 in the first layer gave similar results. We also directly compared predictive learning rule with backpropagation through time (BPTT) on the same convolutional network (Fig. 3). To ensure generality of this comparison, we repeated the training with BPTT three times using different learning rates for each simulation. Using BPTT with the same learning rates as in our predictive model (0.4, 0.028, 0.025), the error rate was 20.88%. For BPTT with learning rate of 0.1 for all layers, the error rate was 21.23%, and 22.77% for a learning rate of 0.2 (Fig. 3B). Code for convolutional network was adopted from (*79*), which we modified to include our predictive learning rule. To reproduce our results, our code for the convolutional network with all implementation details is available at: https://github.com/ykubo82/bioCHL/tree/master/conv. Altogether, those results show that our predictive learning rule can also be successfully implemented in deeper networks and on more challenging tasks.

### Surgery, recording and neuronal data

The experimental procedures for the awake, head-fixed experiment have been previously described (*42, 43*) and were approved by the Rutgers University Animal Care and Use Committee, and conformed to NIH Guidelines on the Care and Use of Laboratory Animals. Briefly, a headpost was implanted on the skull of four Sprague-Dawley male rats (300-500g) under ketamine-xylazine anesthesia, and a craniotomy was performed above the auditory cortex and covered with wax and dental acrylic. After recovery the animal was trained for 6-8 days to remain motionless in the restraining apparatus. On the day of the surgery, the animal was briefly anesthetized with isoflurane, the dura was resected, and after a recovery period, recording began. For recording we used silicon microelectrodes (Neuronexus technologies, Ann Arbor MI) consisting of 8 or 4 shanks spaced by 200µm, with a tetrode recording configuration on each shank. Electrodes were inserted in layer V in the primary auditory cortex. Units were isolated by a semiautomatic algorithm (klustakwik.sourceforge.net) followed by manual clustering (klusters.sourceforge.net)(*80*). Only neurons with average stimulus evoked firing rates higher than 3 SD above pre-stimulus baseline were used in analysis, resulting in 9, 12, 12, and 22 neurons from each rat. For predicting evoked activity from spontaneous, we also required that neurons must have mean firing rate during spontaneous packets above said threshold which reduced the number of neurons to 40. The spontaneous packet onsets were identified from the spiking activity of all recorded cells as the time of the first spike marking a transition from a period of global silence (30 ms with at most one spike from any cell) to a period of activity (60 ms with at least 15 spikes from any cells), as described before in (*42, 75*).

## Acknowledgments

This work was supported by Compute Canada, NSERC and CIHR grants to AL, and DARPA #HR0011-18-2-0021 and NIH #MH125557 to BLM. We thank Karim Ali, Lukas Grasse, Mikhail Klassen and Reza Torabi for help, and we thank Peter Bartho for sharing data.

## Author contributions

AL conceived the project, analyzed data, performed computer simulations and wrote the manuscript; BLM engaged in theoretical discussions and commented extensively on the manuscript. YK performed computer simulations and contributed to writing manuscript.

## Data sharing

Our code is publicly available at: https://github.com/ykubo82/bioCHL. The data presented here are available from the corresponding author upon request.

## Suppl. Materials

**Suppl. Fig. 1.**
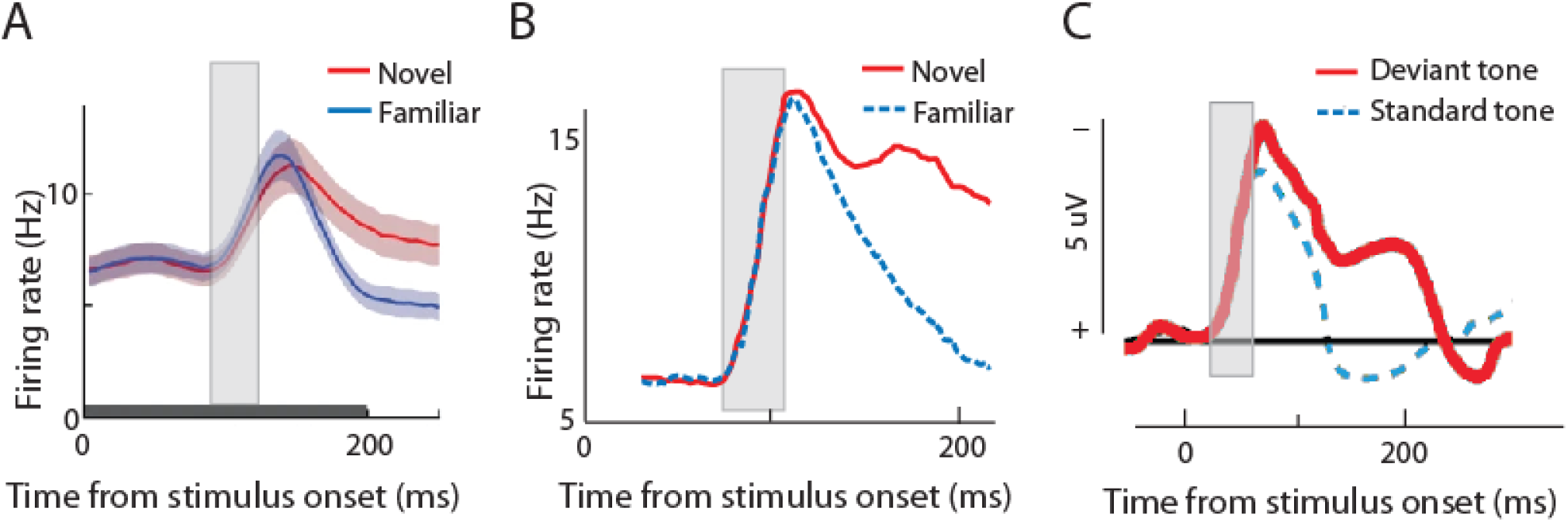
Consistently with our model, experiments in humans and monkeys show that activity corresponding to ‘free phase’ is followed by equivalent of ‘clamped phase’. (A) Average population activity across 73 inferior temporal cortex excitatory neurons in monkeys during passive viewing of novel (red) and familiar stimuli (blue; adopted from Lim et al. 2015 after Woloszyn et al. 2012). Note that consistently with our model, for both types of stimuli the neuronal response are initially the same (denoted by gray area). However, later response to novel stimuli diverges from the familiar, likely due to top-down modulation from other brain regions. In the context of our work, response to familiar stimuli could be seen as expected activity equivalent to ‘free phase’, and response to novel stimuli could be interpreted as activity with additional top-down modulation analogous to ‘clamped phase’ in our model. Our learning rule suggests that the late response to novel stimuli which deviates from expected activity can provide training signal for synaptic update. (B) Similar results from a different lab. Average population activity across 298 inferior temporal cortex neurons in monkeys during passive viewing of novel (red line) and familiar stimuli (dashed blue line; adopted from Freedman et. al, 2006). (C) Average event-related potential recorded with EEG electrode located above the frontal cortex in healthy human adults in response to 1000 Hz standard tones and to 1032 Hz deviant tones (20% probability; adopted from Sams et. al 1985). This is a typical example from a rich body of scientific literature on the mismatch negativity phenomenon (a brain response to violations of expected stimulus rule). The similarity of neuronal responses at early phase and divergence from expected response for unexpected stimuli only in the later phase provides another biological example as justification for combining free and clamped phase in our model.

**Suppl. Fig. 2.**
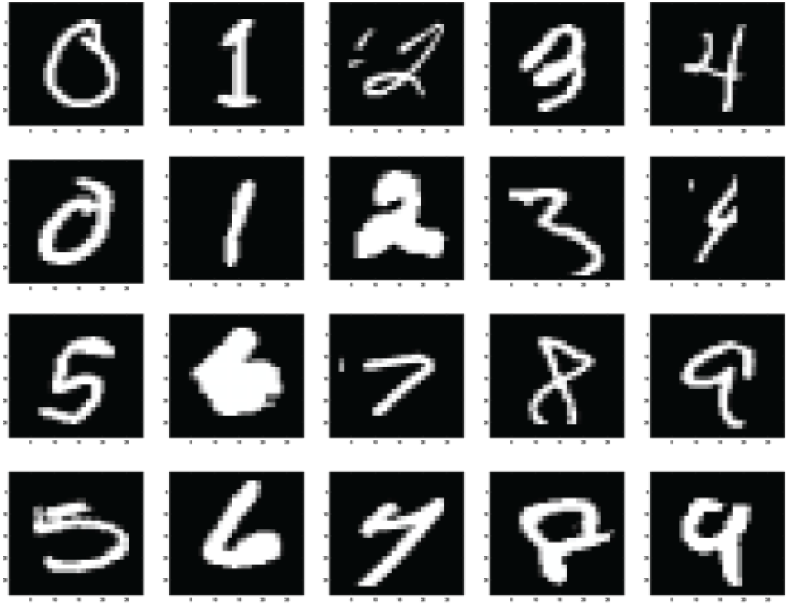
Examples of handwritten digits from MNIST data set (LeCun et al. 1998). Note that classifying such images is a non-trivial task even for humans, as for instance digits at the bottom row (5, 6, 7, 8, 9) could be mistaken for: 0, 4, 4, 9 and 4, respectively.

**Suppl. Fig. 3.**
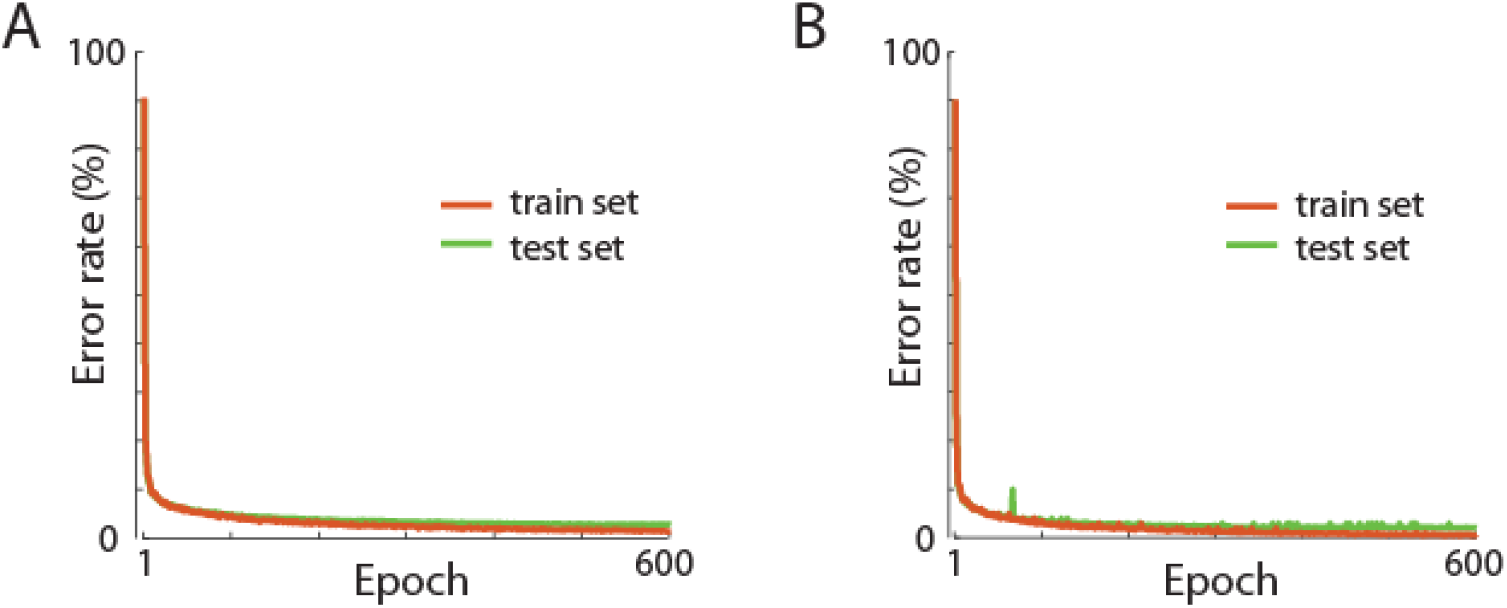
(A) The learning curve for a network with 80% of excitatory and 20% inhibitory neurons in the hidden layer. For each excitatory neuron, all outgoing synaptic connections to output neurons were non-negative (w ≥ 0), and for inhibitory neurons all outgoing connections to output neurons were negative or zero (w≤0). The network had the same architecture as our main network (784-1000-10 neurons), and was using the same learning rule (Eq. 3). On the test data set it achieved a comparable error rate of 2.66%. Those results are consistent with other models which showed that implementing Dale’s law did not reduce network performance substantially (i.e.: Cornford et al. 2020). (B) Learning curves for a network with asymmetric connections. The network had the same architecture as our main network (784-1000-10 neurons), and on the same MNIST dataset it achieved an error rate of 1.96%. Typical artificial neural networks trained with the backpropagation algorithm, require that a synaptic weight from neuron *i* to *j* (*w*_*ij*_) is exactly the same as a synaptic weight from neuron *j* to *i* (*w*_*ij*_*=w*_*ji*_ is called symmetric weight). However, it was recently shown that networks without symmetric connections can still converge on a good solution (Lillicrap et al. 2016, Akrout et al. 2019). Similar results were also found by Detorakis et al. (2019) who used asymmetric weights with Contrastive Hebbian Learning. Thus, our results are consistent with that previous work and show that our learning rule also works in such more biologically plausible network configurations.

**Suppl. Fig. 4.**
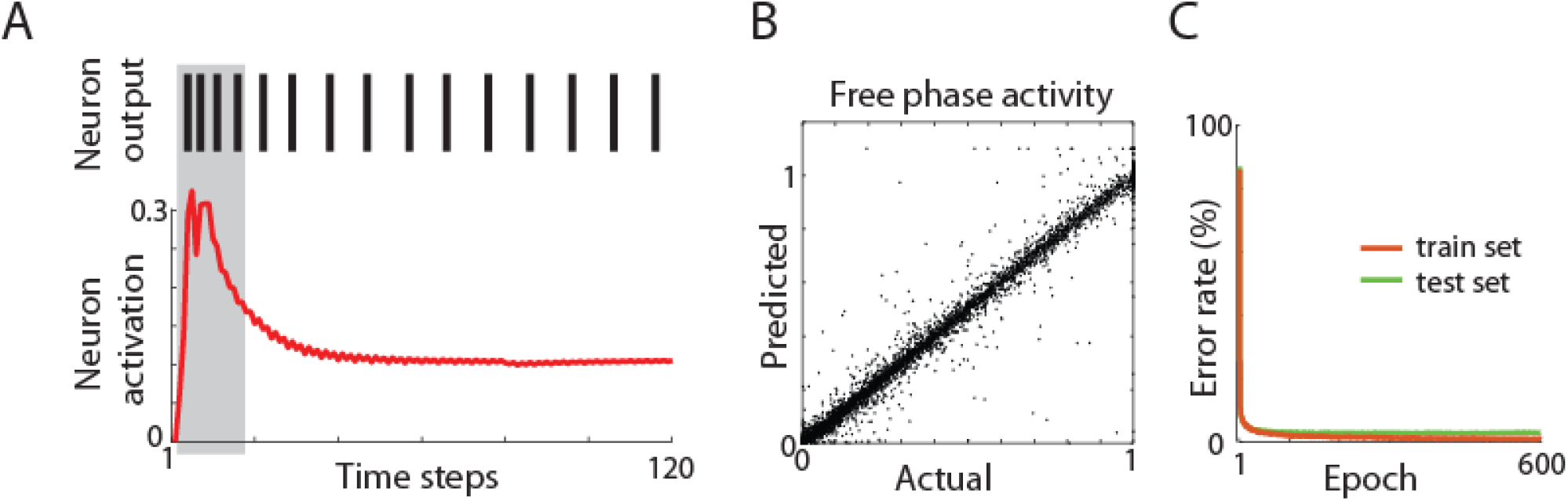
Implementation of predictive learning rule in a spiking neuronal network. (A) Activity of a sample neuron. Red trace shows neuron internal activation which is a function of synaptic inputs. (Top) Spikes are generated based on the value of internal activation. Gray shaded area marks the extent of free phase which is used to predict steady-state activation. (B) Actual vs. predicted steady-state free phase activity. Each dot represents the activity of one neuron during presentation of one stimuli. Activity of 1000 hidden neurons during presentation of 200 stimuli from the test dataset is shown. The correlation coefficient between actual and predicted activity was R=0.99 (p<0.00001). (C) Learning curves on MNIST task for training and testing dataset. Error rate of 2.46% on the test set shows that the spiking neuronal network with our learning rule was able to solve the presented task.

### Spiking network model

The code for this network with all implementation details is provided at: https://github.com/ykubo82/bioCHL/tree/master/bioCHLspk. Briefly, our network was based on work by O’Connor et al. (2019) who developed spiking networks using the Equilibrium Propagation algorithm which is an expansion of Contrastive Hebbian Learning (Scellier et al., 2017). The steady state is calculated using the Forward Euler method with each time step 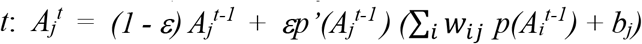, where *A* is the activation state, *p*’(*A*_*j*_) is the derivative of the activation function *p, b* is the bias, and *i* and *j* are neurons indices. *ε* can be seen as the learning rate of the activations. These states are clipped to range [0, 1]. Each neuron communicates only using binary signals: 0 or 1, to model spiking activity. For generating spikes, an encoder converts the neuron activation state to 1 or 0, which is sent as an output signal (panel A). The binary signals received by a neuron are converted into the activation state by a decoder. This could be seen as converting discrete spikes into a continuous post-synaptic membrane potential in actual neurons. We modified this spiking network by implementing our learning rule and by adding a least-squares model for predicting steady-state activation based on initial activation in steps 1-17. We also used AdaGrad (Duchi et al., 2011) to search for hyperparameters, which resulted in setting the learning rate to 0.01. The network architecture was the same as in our main model with 784-1000-10 neurons. Those results are consistent with other studies which showed that spiking networks can perform as well as non-spiking networks (i.e.: Zenke & Ganguli, 2018; Bellec et al. 2020).

### Acoustic stimuli

In animal experiments, as stimuli we used 1 s long pure tones (2, 3.3, 5.4, 8.9, 15 and 24 kHz at 60dB), interleaved with 1s periods of silence, as described in (Luczak et al. 2009 & 2013). Activity occurring >200ms after stimulus offset and before the next stimulus onset was regarded as spontaneous. The stimuli were continuously presented for 20-108 min, depending on how long the animal appeared to be comfortable during the experiment. All stimuli were tapered at the beginning and end with a 5ms cosine window. Experiments took place in a single-walled sound isolation chamber (IAC, Bronx, NY) with sounds presented free field (RP2/ES1, Tucker-Davis, Alachua, FL). To compensate for the transfer function of the acoustic chamber, tone amplitudes were calibrated prior to the experiment using a condenser microphone placed next to the animal’s head (7017, ACO Pacific, Belmont CA) and a MA3 microphone amplifier (Tucker-Davis).

### Changes in neuronal activity across periods

For each animal we divided the duration of recording into two halves. For each neuron we calculated the mean stimulus evoked firing rate in 30-40 ms time window, averaged over all stimulus presentations in each half of the experiment. Similarly, for each neuron we calculated the average predicted activity for the 30-40 ms time window, averaged across all stimuli in the 1st half of the experiment. We then calculated the difference between activities in the 2nd half minus the 1st half, and we correlated it with the difference between activity in the 1st half minus predicted activity in the 1st half (Fig. 3D). As described in the Methods section, only neurons with average stimulus evoked firing rates higher than 3 SD above pre-stimulus baseline were used in our analyses. Dividing the data in different proportions e.g. 60%-40% gave the same conclusions.

For comparison, analogous analyses were done on artificial neurons (Fig. 3C). For each neuron in the hidden layer the steady-state clamped activity was averaged over all 1200 stimulus presentations in a single learning epoch. Similarly, for each neuron in the hidden layer, the predicted activity was averaged across all stimuli presented in an epoch. We then calculated the difference between clamped activities in 2 epoch: 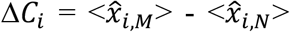, where <> denotes average, *i* is index of neuron, and *M* and *N* are indexes of epochs with *M* > *N*. This was then correlated with difference between average clamped and predicted activity in epoch *N*: 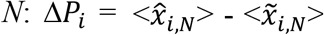. We found that the correlation between Δ*C* and Δ*P* was strongest in the earliest epochs where the learning curve was most changing (Fig. 2E). For this simulation we used neural network as described in Methods with 784-1000-10 neurons, and we used the learning rule described in Eq. 3. However, in order to reproduce the repeated presentation of the same stimuli as in animal experiments, we used the same 1200 stimulus in each learning epoch. Presenting different stimuli at each epoch as in our original simulation, gave qualitatively similar results.

### Predicting evoked dynamics from spontaneous activity

For predicting evoked dynamics from spontaneous, first we analyzed ‘types’ of spontaneous activity patterns. For this we divided spontaneous packets into 10 groups based on principal component analysis (PCA). Specifically, each packet was represented as a NxT matrix, where N is number of neurons and T=30 is number of 3ms long time bins. This matrix was then converted into a single vector V with length of NxT. This was repeated for each packet, thus we obtained a VxP matrix, where P is the number of spontaneous packets (P for each animal was: 293, 855, 1215 and 2472). We then ran PCA on the VxP matrix and based on values of the 1st PC we divided packets into 10 groups. Note that the distribution of packets in PCA space was continuous, thus from this analysis it should not be interpreted that there are distinct types of packets. Dividing packets into 5-20 groups and including the 2nd and 3rd PC gave consistent results with those presented in the main text.

For predicting evoked dynamics from spontaneous, we faced the problem that defining the onset of spontaneous packets is less accurate than the onset of evoked responses. Thus, to ensure a similar precision of alignment, during each stimulus presentation we detected the onset of evoked responses using the same criteria as for spontaneous packets. Then, the evoked responses were realigned according to that of the detected onset. This allowed for estimating the spontaneous and evoked activity with the same precision. Using the original onset times instead of the realigned gave qualitatively similar results.

**Suppl. Fig. 5.**
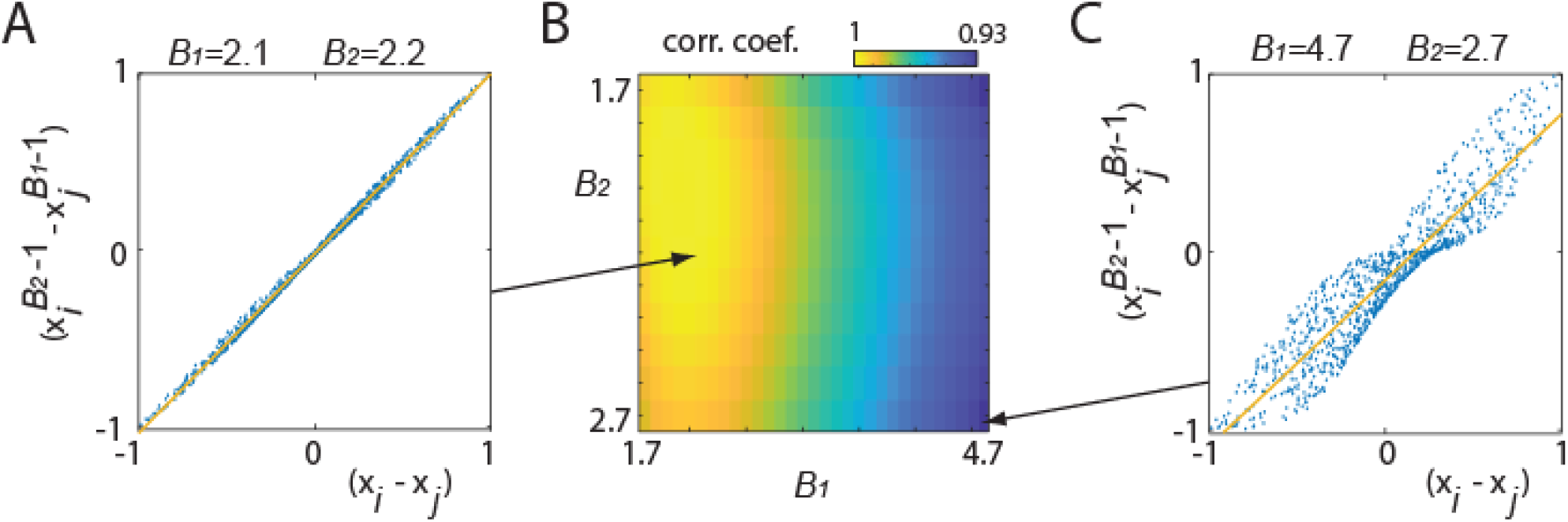
Testing effect of different values of *β*_*1*_ and *β*_*2*_. In Eq. 6 we used: *β*_*1*_ = 2 and *β*_*2*_ = 2, which allowed to reduce that formula from exponential to linear. Here we illustrate that resulted linearized expression in form: (*x*_*i*_ − *x*_*j*_) is a reasonable approximation of non-linear version: 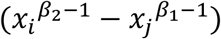 for *β*_*1*_ and *β*_*2*_ in range observed experimentally: 1.7 < *β*_*1*_ < 4.8 and 1.7 < *β*_*2*_ < 2.7 (Devor et al., 2003), and for typical values of *x* in range between 0-1. (A) Relation between (*x*_*i*_ − *x*_*j*_) and 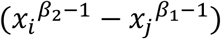 for *β*_*1*_ = 2.1 and *β*_*2*_ = 2.2. Here we generated 1000 random values for *x*_*i*_ and separately for *x*_*j*_, uniformly distributed on interval 0-1. Distribution of points along diagonal shows close to linear relation between results of both formulas. Least square regression line is shown in yellow. Correlation coefficient between points obtained with linear and non-linear formula was R=0.9988. (B) Similarly, we calculated correlation coefficients between both formulas for all combinations of *β*_*1*_ and *β*_*2*_ in range: 1.7 ≤ *β*_*1*_ ≤ 4.8 and 1.7 ≤ *β*_*2*_ ≤ 2.7, using step size of 0.1. The lowest correlation coefficient was R = 0.9335 for *β*_*1*_ = 4.7 and *β*_*2*_ = 2.7. (C) The same plot as in panel A, but for the “worst case” scenario with *β*_*1*_ = 4.7 and *β*_*2*_ = 2.7. Note that even in this case, the relation between both formulas is close to linear.

### Maximizing future energy balance

Intuitively, it makes sense that planning, i.e. making predictions, can improve success of an organisms in accessing more energy resources. In this section we show that this holds true even for a single neuron, where maximizing future energy balance is best achieved by predicting future activity. For that, first we will write equation for energy balance for a neuron at time *t+n*, where *t* represents current time and *n* is a small time increment. Using the same logic and notation as in Eq. 4, energy balance of a neuron *j* at time *t+n*, can be expressed as a function of: cost of housekeeping processes (*ε*), cost of electrical activity (which for simplified linear neurons can be written as sum of its synaptic inputs: *x*_*j,t*+*n*_ = ∑_*i*_ *w*_*ij*_*x*_*i,t*+*n*_), and energy supply from local blood vessels controlled by combined local activity of neurons (∑_*k*_ *x*_*k,t*+*n*_):

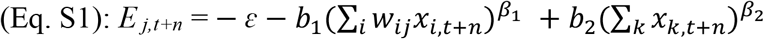

where *β*_*1*_ and *β*_*2*_ describe a non-linear relation between activity and energy (Devor et al., 2003).

In the main text we show in simulations (Fig. 2) and in experimental data (Fig. 3), that for small *n*, activity of neuron *j* at time *t+n* could be approximated by a linear function of its activity at earlier time step *t*: *x*_*j,t*+*n*_ = *λ*_*j*_*x*_*j,t*_, where *λ* is a regression coefficient. Thus Eq. S1 can be rewritten as

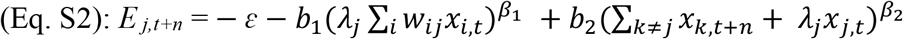

Using gradient ascent method, we can calculate change in weights to maximize future energy balance:

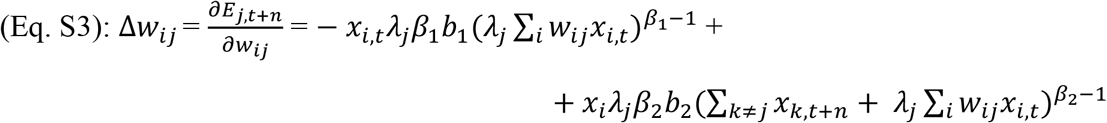

Note that in Eq. S3: 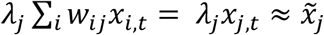, thus this term corresponds to predicted future activity: 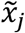. We will also denote a population activity ∑_*k*≠*j*_ *x*_*k,t*+*n*_ as: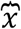, which simplifies Eq. S3 to:

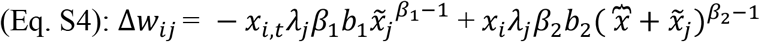

after factoring out *x*_*i,t*_*λ*_*j*_ and switching order of terms, we get:

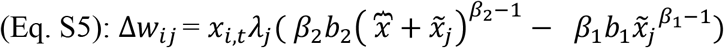

Considering results from Suppl. Fig. 6, we can use *β*_*1*_ = 2 and *β*_*2*_ = 2, which allows to simplify and reorganize terms as follows:

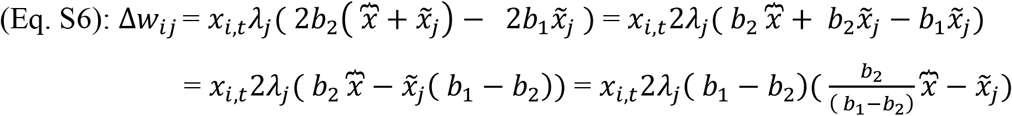

after denoting constant terms as α_3_ = 2*λ*_*j*_(*b*_1_ − *b*_2_), and as 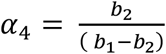, we obtain:

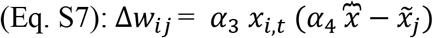

This shows that maximizing future energy balance requires neuron to predict its future activity: 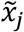. Also note that in the brain, networks are highly recurrent, thus population activity 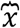 can be seen as providing similar role as top-down modulation: 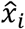. Altogether, this suggests, that type of a learning rule as in Eq. 3: 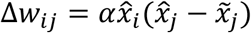, may be a necessity to allow neuron for its highly energy intensive operation.

Note also that this derivation of learning rule from metabolic principles is obtained simply by only considering input and output energy to the cell, without need for including specific metabolic interactions. This provides an important theoretical contribution by showing how complex relations between cell metabolism and plasticity could be described more simply.

### Activation function

To simplify presented derivation of learning rule we used linear model of a neuron. However, we can obtain the same expression even if we use non-linear neuronal model with activation function like ReLU: f(x) = x^+^ = max(0, x) (Glorot et al. 2011). This is because for *x* ≥ 0: ReLU(*x*) = *x*, which results in the same formulas as presented in our derivation. The only difference is that if we use ReLU then we need to write a separate expression for *x* < 0, to specify that in such case ReLU(*x*) = 0. For sigmoid activation function, showing equivalence of the formulas could be more complicated. However, there are multiple arguments that ReLU could be a better model of biological neurons than logistic sigmoid neurons (Glorot et al. 2011), and our neural network simulations performed similarly for both neuron models.

